# Dopamine and ALK4 signaling synergize to induce PCBP1-mediated alternative splicing of FosB and sustained behavioral sensitization to cocaine

**DOI:** 10.1101/2022.01.20.477040

**Authors:** Favio A. Krapacher, Diana Fernández-Suárez, Annika Andersson, Alvaro Carrier-Ruiz, Carlos F. Ibáñez

## Abstract

ΔFosB, an alternative spliced product of FosB, is an essential component of dopamine-induced reward pathways and a master switch for addiction. However, the molecular mechanisms of its generation and regulation by dopamine signaling are unknown. Here we report that dopamine D1 receptor signaling synergizes with the activin/ALK4/Smad3 pathway to potentiate the generation of ΔFosB mRNA in medium spiny neurons (MSNs) of the nucleus-accumbens (NAc) through activation of the RNA binding protein PCBP1, a regulator of mRNA splicing. Concurrent activation of PCBP1 and Smad3 by D1 and ALK4 signaling induced their interaction, nuclear translocation, and binding to sequences in exon-4 and intron-4 of FosB mRNA. Ablation of either ALK4 or PCBP1 in MSNs impaired ΔFosB mRNA induction and nuclear translocation of ΔFosB protein in response to repeated co-stimulation of D1 and ALK4 receptors. Importantly, ALK4 was required in NAc MSNs of adult mice for behavioral sensitization to cocaine. These findings uncover an unexpected mechanism for ΔFosB generation and drug-induced sensitization through convergent dopamine and ALK4 signaling.

## Introduction

The central dopamine system of the mammalian brain, notably the ventral tegmental area (VTA), substantia nigra pars compacta (SNc) and their target neurons in the nucleus accumbens (NAc), has a pivotal role in the regulation of responses to natural rewards such as food, drink, sex and social interaction (Kauer and Malenka, 2007; Volkow and Morales, 2015). One of its main functions is to reinforce behaviors that lead to successful earning of such rewards to the extent they enhance survival and reproduction. On the flip side, this also makes the dopamine system a vulnerable target that can be hijacked by a variety of psychoactive stimuli, including drugs of abuse, as well as compulsive behaviors, such as pathological overeating, compulsive running and gambling (Lüscher, 2016). An important area of neuroscience research is centered on understanding the intricate ways in which drugs of abuse alter the brain to produce the functional and behavioral abnormalities that characterize addiction. The persistence of these alterations is thought to be sustained by transcriptional changes that maintain long lasting molecular and structural adaptations in neuronal circuits involved in reward processing and its behavioral manifestation (Teague and Nestler, 2021). A key component of transcriptional mechanisms underlying reward responses is ΔFosB, a transcription factor encoded by a splice variant of FosB, a member of the Fos family of immediate early genes (Nestler, 2008; 2013; Ruffle, 2014). Fos family proteins are rapidly induced in the NAc after acute exposure to many drugs of abuse, such as cocaine, but their levels return to baseline within hours of drug administration due to protein instability (Graybiel et al., 1990; Hope et al., 1992; Young et al., 1991). Upon chronic drug exposure, most Fos family proteins show desensitization and fail to accumulate to significant levels (Alibhai et al., 2007). In contrast, repeated drug administration results in the accumulation of ΔFosB which, due to its unusually high stability, can persist for days or weeks in NAc neurons (Chen et al., 1997; 1995; Hiroi et al., 1997). ΔFosB mRNA is generated by the removal of intron 4 (also known as the regulated intron) from the 4^th^ and last exon of FosB mRNA, thereby introducing a frameshift that leads to premature termination of the protein. As a consequence, ΔFosB lacks degradation signals present in the C-terminus of FosB. In addition, various phosphorylation events at the N-terminus of ΔFosB also contribute to its stabilization (Nestler, 2013). The stability of ΔFosB has been viewed as an example of the ways in which drug-induced alterations in gene expression can persist for long periods of time after drug withdrawal. Longer term changes in gene transcription can be further propagated by the ability of ΔFosB to induce histone modifications in target genes (Teague and Nestler, 2021). Functional studies in transgenic mice using inducible overexpression of ΔFosB or AP1 dominant negative proteins in medium spiny neurons (MSNs) of the NAc support the notion that accumulation of ΔFosB increases sensitivity to the locomotor and rewarding effects of cocaine and other drugs of abuse, increases cocaine self-administration, and promotes positive reinforcement (Kelz et al., 1999; Peakman et al., 2003; Zachariou et al., 2006). Based on these results, it has been proposed that ΔFosB serves as a critical molecular switch for addiction (Nestler et al., 2001; Nestler, 2013).

Despite the importance of ΔFosB as a key regulator of reward responses, very little is known about the molecular mechanisms that mediate its production by alternative splicing of FosB mRNA. The RNA binding protein PTB (polypyrimidine tract binding protein) was found to interact with the 3’ end of intron 4 of FosB mRNA, and PTB overexpression in HeLa or PC12 cells decreased the splicing efficiency of a FosB minigene transcript, favoring retention of intron 4 (Alibhai et al., 2007; Marinescu et al., 2007). These results suggested that PTB may be a negative regulator of ΔFosB production. However, the actual role of this protein in ΔFosB induction is unclear, as PTB was not regulated by either acute or chronic drug administration (Alibhai et al., 2007), and splicing of FosB mRNA intron 4 was not affected by PTB knockdown (Marinescu et al., 2007). Based on these and other findings, it has been postulated that the generation of ΔFosB mRNA is constitutive and essentially proportional to the levels of FosB mRNA, specially under conditions of high gene induction (Alibhai et al., 2007). Accumulation of ΔFosB in NAc during chronic drug exposure is thought to be solely due to the high stability of the protein, not to increased generation of ΔFosB mRNA, and evidence that dopamine signaling can regulate FosB mRNA splicing to contribute to the generation of ΔFosB is currently lacking.

Activin receptor-like kinase 4 (ALK4) is a type I serine-threonine kinase receptor for a subset of TGFβ superfamily ligands that includes activins A, B and members of the Growth Differentiation Factor (GDF) subfamily, such as GDF-1, -3 and -5 (Dijke et al., 1994; Moustakas and Heldin, 2009; Schmierer and Hill, 2007). ALK4 is an essential component of activin A signaling, while activin B, GDF-1 and GDF-3 can signal by either ALK4 or the related receptor ALK7 (Andersson et al., 2008; 2006; Tsuchida et al., 2004). The main downstream effectors of activated TGFβ superfamily receptors are the Smad proteins (Derynck and Budi, 2019). Canonical signaling by ALK4 involves the recruitment and phosphorylation of Smad2 and Smad3, which then partner with Smad4 and translocate to the cell nucleus, where they regulate gene expression in conjunction with cell type-specific transcription factors (Budi et al., 2017; Massagué, 2012; Schmierer and Hill, 2007). ALK4 is expressed in the extra-embryonic ectoderm and epiblast of early post-implantation embryos; by mid-gestation, its expression is nearly ubiquitous, including the central nervous system (Gu et al., 1998; Verschueren et al., 1995). A null mutation in mouse *Acvr1b*, the gene encoding ALK4, is embryonic lethal due to defects in primitive streak formation and gastrulation (Gu et al., 1998). Using a conditional approach, a recent study from our laboratory demonstrated a role for ALK4 in regulation of cortical somatostatin interneuron development (Göngrich et al., 2020). However, a large gap of knowledge remains on the functions of ALK4 in the nervous system, particularly in the adult brain. Various activities have been attributed to activins and related ligands in nervous tissue, specially in paradigms of neural injury and morphological plasticity, mainly through gain-of-function studies using exogenous activin administration (Hasegawa et al., 2014; Su et al., 2018). In the context of drug-induced behaviors, it has been reported that infusion of activin A in the NAc increases the rate of cocaine-self administration in rats (Gancarz et al., 2015). However, the role, if any, of endogenous activin/ALK4 signaling in the NAc, or its possible relationship to molecular mechanisms of reward, such as ΔFosB, remain unknown.

In this study, we investigated the contribution of mechanisms operating at the mRNA level to the regulation of ΔFosB production after stimulation of dopamine D1 receptors in primary cultures of NAc MSNs using a novel *in vitro* sensitization paradigm. We discovered a previously unappreciated synergy between dopamine and activin/ALK4 signaling leading to robust generation of ΔFosB mRNA. Further exploration of this finding led to the identification of the RNA binding protein PCBP1 as the first known positive regulator of FosB mRNA splicing, and validation of its requirement for ΔFosB mRNA generation in response to concurrent dopamine and activin signaling. Strikingly, ablation of ALK4 in NAc MSNs of adult mice completely abolished drug-induced induction of ΔFosB mRNA and behavioral sensitization to cocaine.

## Results

### Dopamine and ALK4 signaling synergize to potentiate induction of ΔFosB mRNA and nuclear translocation of ΔFosB protein in NAc MSNs

Sensitization is the phenomenon by which exposure to a stimulus induces a larger response to a subsequent exposure. Studies in rats and mice have shown that a single exposure to a psychostimulant, such as cocaine, can induce behavioral sensitization in both a context-dependent and a context-independent manner. Since enhanced responses to the second exposure can be observed days or even weeks after the first, this paradigm has been utilized to study persistent neural plasticity induced by psychoactive drugs and other stimuli. We investigated the effects of stimulation of dopamine receptors in primary cultures of NAc MSNs using an *in vitro* sensitization paradigm consisting of two consecutive treatments with the D1 agonist SKF81297 (Fig. 1A). A single 2h stimulation of MSNs with SKF81297 induced a small (1.7-fold) increase in FosB mRNA levels, but no significant change could be observed in c-Fos or ΔFosB mRNAs, nor in the levels of ARC and EGR1 (a.k.a. Ziff268) mRNAs, two other critical mediators of transcriptional responses to cocaine and D1 signaling (Jiang et al., 2021; Penrod et al., 2020; Renthal et al., 2009; Valjent et al., 2006) (Fig. S1A-F). Likewise, nuclear translocation of ΔFosB protein was also unaltered by a single stimulation with SKF81297 (Fig. S1G, H). In contrast, two consecutive 2h treatments separated by 2 days produced a 3-fold increase in ΔFosB mRNA (Fig. 1A, B). Induction of FosB mRNA remained at about 2-fold (Fig. 1C), while no significant changes were observed in c-Fos, ARC and EGR1 mRNAs (Fig 1D-F). Stimulation of ALK4 receptors with activin A had no effect on ΔFosB mRNA levels, either after a single treatment or two consecutive exposures (Fig. 1B, S1B). Neither were c-Fos, FosB, ARC or EGR1 mRNA levels affected by activin A on its own (Fig. 1C-F, S1C, D, G, H). On the other hand, the response of ΔFosB mRNA to two consecutive treatments with SKF81297 was significantly augmented (>10-fold) by co-stimulation with activin A (Fig. 1B). Under these conditions, herein termed consecutive co-stimulation, FosB and ARC mRNAs levels increased 3-fold, and EGR1 mRNA 6-fold, while no change was observed in c-Fos mRNA (Fig. 1C-F). Importantly, the response of ΔFosB mRNA to consecutive co-stimulation with SKF81297 and activin A was paralleled by induction of ΔFosB protein and its translocation to MSNs nuclei (Fig. 1G, H). Although activin A had no effect on its own, it greatly potentiated the effect of SKF81297 on the nuclear translocation of ΔFosB protein (Fig. 1H). Neither SKF81297, activin A or its combination affected the levels of Acvr1b mRNA, encoding ALK4, in MSN primary cultures (Fig. S1I).

**Figure 1.**
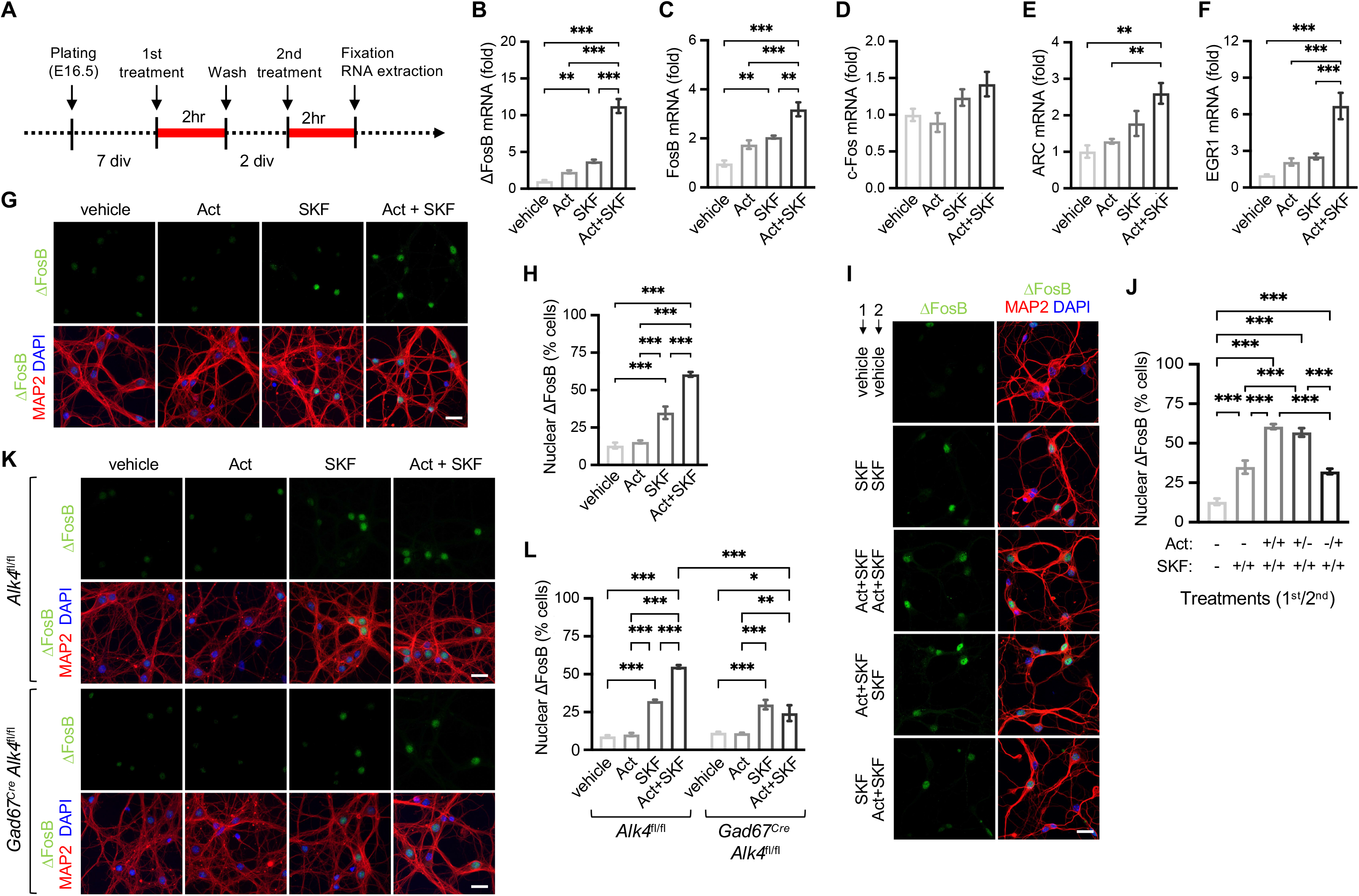
Dopamine and ALK4 signaling synergize to potentiate induction of ΔFosB mRNA and nuclear translocation of ΔFosB protein in NAc MSNs. (A) Schematic of the protocol used for consecutive stimulation of primary cultures of NAc MSNs. MSNs were extracted at embryonic day 16.5 (E16.5) and cultured for 7 days (7div) before 2 hours (2hr) treatment with either 1μM SKF81297 (SKF), 100ng/ml activin A (Act) or their combination (Act+SKF). Following 2 additional days (2div), the treatments were repeated and cultures fixed for imaging or collected for RNA extraction. (B-F) Real-time quantitative PCR (RT-qPCR) analysis of ΔFosB (B), FosB (C), c-Fos (D), ARC (E) and EGR1 (F) mRNAs in MSNs cultures exposed to consecutive stimulation (same treatment in both treatment days) with activin A (Act), SKF81297 (SKF) or their combination as indicated. Results are expressed as average ± SEM of fold increase over vehicle . **, p<0.01; ***, p<0.001; N=5; one-way ANOVA followed with Tukey’s multiple comparisons test. (G) Representative immunofluorescence staining of ΔFosB (green) in cultured MSNs exposed to consecutive stimulation (same treatment in both treatment days) with activin A (Act), SKF81297 (SKF) or their combination as indicated. Counterstaining with MAP2 and DAPI is shown in the lower row. Scale bar, 25μM. (H) Quantification of ΔFosB^+^ nuclei in cultured MSNs after the indicated treatments (same treatment in both treatment days). Results are expressed as average ± SEM of % of MAP2^+^ cells displaying ΔFosB in the nucleus. ***, p<0.001; N=6; one-way ANOVA followed with Tukey’s multiple comparisons test. (I) Representative immunofluorescence staining of ΔFosB (green) in cultured MSNs exposed to consecutive stimulation, with different 1^st^ and 2^nd^ treatments with activin A (Act), SKF81297 (SKF) or their combination as indicated. Counterstaining with MAP2 and DAPI is shown in the right column. Scale bar, 25μM. (J) Quantification of ΔFosB^+^ nuclei in cultured MSNs after different 1^st^ and 2^nd^ treatments with activin A (Act), SKF81297 (SKF) or their combination as indicated. Results are expressed as average ± SEM of % of MAP2^+^ cells displaying ΔFosB in the nucleus. ***, p<0.001; N=6; one-way ANOVA followed with Tukey’s multiple comparisons test. (K) Representative immunofluorescence staining of ΔFosB (green) in cultured MSNs extracted from mutant mice lacking ALK4 in GABAergic neurons (*Gad67*^Cre^;*Alk4*^fl/fl^) and control mice (*Alk4*^fl/fl^) after exposure to consecutive stimulation (same treatment in both treatment days) with activin A (Act), SKF81297 (SKF) or their combination as indicated. Counterstaining with MAP2 and DAPI is shown in the second and fourth rows. Scale bar, 25μM. (L) Quantification of ΔFosB^+^ nuclei in cultured MSNs from mutant and control mice after the indicated treatments (same treatment in both treatment days). Results are expressed as average ± SEM of % of MAP2^+^ cells displaying ΔFosB in the nucleus. *, p<0.05; **, p<0.01;***, p<0.001; N=6; one-way ANOVA followed with Tukey’s multiple comparisons test.

The changes in responsiveness that accompany sensitization to a second exposure to a psychostimulant are thought to be induced by the first stimulation (“induction phase”), while the second stimulation serves to reveal the underlying sensitization effect (“expression phase”). The strong synergy between SKF81297 and activin A after consecutive co-stimulation prompted us to investigate whether activin A exerted its potentiation effects during the first exposure, the second or both. In order to test this, activin A was provided along with SKF81297 only during the first (“induction”) or second (“expression”) stimulation of MSN primary cultures. These were compared to consecutive treatments with SKF81297 alone or together with activin A. Only when administered along with SKF81297 in the 1^st^ treatment (induction phase) could activin A potentiate nuclear translocation of ΔFosB protein in MSNs, fully recapitulating the effect of consecutive co-stimulation (Fig. 1I, J). No potentiation was observed if activin A was only administered in the 2^nd^ treatment (expression phase), indicating that the potentiation effects of activin A on dopamine receptor signaling take place during the induction phase of sensitization. The effects of two consecutive co-stimulations with SKF81297 and activin A in ΔFosB mRNA induction and ΔFosB protein nuclear translocation were significantly reduced by SB431542, an inhibitor of ALK4, the main activin receptor, and PKI 14-22, an inhibitor of Protein Kinase A (PKA), a downstream effector of D1 receptor signaling, respectively (Fig. S1J-L).

Acvr1b mRNA, encoding ALK4, was found to be expressed in 70 to 75% of DARPP-32^+^ and D1^+^ neurons in the NAc and striatum (Str) (Fig. S2A-C). These neurons are GABAergic, and *Gad67*^Cre^;*Alk4*^fl/fl^ mutant mice expressing Cre recombinase in GABAergic neurons showed a significant reduction in Acvr1b mRNA levels in the NAc and Str, but less so in other areas, including motor cortex (Fig. S2D-F). ALK4 protein expression, as detected by immunohistochemistry, was also reduced in NAc, but not in primary motor cortex, of these mutant mice (Fig. S2G, H). Nuclear translocation of ΔFosB protein induced by two consecutive co-stimulations with SKF81297 and activin A was investigated in primary cultures of MSNs isolated from these mutant mice. Activin A was unable to potentiate the effects of SKF81297 in MSNs isolated from *Gad67*^Cre^;*Alk4*^fl/fl^ mutant mice (Fig. 1K, L), although these neurons responded normally to SKF81297 alone (Fig. 1L).

### Concurrent signaling by dopamine and ALK4 induces phosphorylation, nuclear translocation, co-localization and direct interaction of Smad3 and the RNA binding protein PCBP1 in NAc MSNs

We considered the possibility that ALK4 signaling may play a role in the generation of ΔFosB mRNA by enhancing the alternative splicing of FosB mRNA. Although Smad proteins were not known to play a role in mRNA splicing, it was shown by Tripathi et al., 2016 that Smad3 could regulate alternative splicing of CD44 mRNA through direct interaction with the RNA-binding protein PCBP1. In addition to Smad3 activation, this process was dependent on concurrent phosphorylation and nuclear translocation of PCBP1 induced by treatment with either epidermal growth factor or serum. The requirement of PKA activity for the synergistic interaction between D1 and ALK4 receptor signaling in ΔFosB mRNA induction and nuclear translocation of ΔFosB protein (Fig. S1J-L) suggested the possibility that PCBP1 may be a target of PKA-mediated phosphorylation. In line with this notion, two consecutive stimulations (as in Fig. 1A) of HeLa cells with forskolin, a strong PKA activator, induced robust phosphorylation of endogenous PCBP1, as detected with antibodies against phospho-PKA substrates, which could be blocked by the PKA inhibitor PKI14-22 (Fig. 2A). Moreover, forskolin induced a strong enrichment of PCBP1 in the nuclear fraction of these cells which could be inhibited by PKI14-22, although it was not significantly affected by activin A or the ALK4 inhibitor SB431542 (Fig. 2B). Smad3 and PCBP1 could be co-immunoprecipitated from lysates of HeLa cells after two consecutive co-stimulations with forskolin and activin A, but not after treatment with either agent alone (Fig. 2C, D). In order to verify these observations in NAc MSNs, we immunostained neuronal cultures for Smad3 and PCBP1 after two consecutive stimulations with SKF81297, activin A or both. As expected, activin A induced robust nuclear translocation of Smad3 in these cultures, while SKF81297 had no effect (Fig. 2E, F). On the other hand, SKF81297 induced nuclear translocation of PCBP1 and co-stimulation with activin A produced a small potentiation of this effect (Fig. 2E, G).

**Figure 2.**
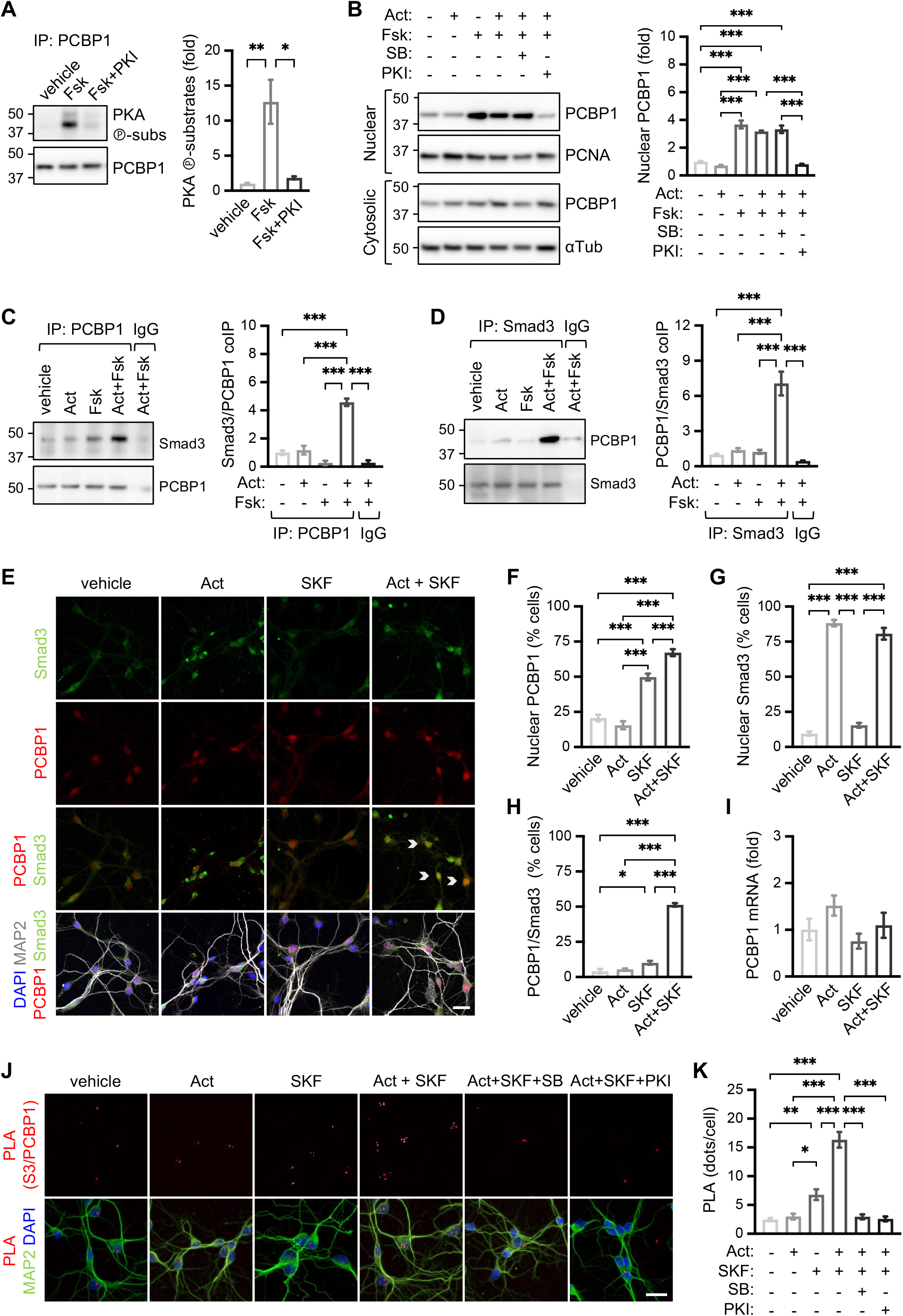
Concurrent signaling by dopamine and ALK4 induce phosphorylation, nuclear translocation, co-localization and direct interaction of Smad3 and the RNA binding protein PCBP1 in NAc MSNs. (A) Western blot analysis of Protein Kinase A phospho-substrates (PKA ℗-subs) in PCBP1 immunoprecipitates from HeLa cells exposed to sequential stimulation (same treatment in both treatment days) with forskolin (Fsk) in the absence or presence of the PKA inhibitor PKI14-22 (PKI). Reprobing with anti-PCBP1 antibodies is shown in the lower blot. Quantification of three independent experiments is shown on the right. Results are expressed as average ± SEM of fold increase over vehicle normalized to PCBP1 levels. *, p<0.05; **, p<0.01, one-way ANOVA followed with Tukey’s multiple comparisons test. (B) Nuclear localization of PCBP1 analyzed by Western blotting of extracts from HeLa cells exposed to sequential stimulation (same treatment in both treatment days) with activin A (Act), forskolin (Fsk), PKA inhibitor (PKI) and the ALK4 inhibitor SB431542 (SB) as indicated. Blots of nuclear and cytosolic fractions were reprobed with anti-PCNA and anti-αTubulin (αTub) antibodies, respectively. Quantification of three independent experiments is shown on the right. Results are expressed as average ± SEM of fold increase over vehicle (first column) normalized to PCBP1 levels. *, p<0.05; **, p<0.01, one-way ANOVA followed with Tukey’s multiple comparisons test. (C) Western blot analysis of Smad3 in PCBP1 immunoprecipitates from HeLa cells exposed to consecutive stimulation (same treatment in both treatment days) with activin A (Act), forskolin (Fsk) or their combination as indicated. The Act+Fsk extracts were also immuoprecipitated using control IgG antibodies as additional control. Reprobing with anti-PCBP1 antibodies is shown in the lower blot. Quantification of four independent experiments is shown on the right. Results are expressed as average ± SEM of fold increase of Smad3/PCBP1 co-immunoprecipitation (coIP) over vehicle (first column) normalized to PCBP1 levels. **, p<0.01, one-way ANOVA followed with Tukey’s multiple comparisons test. (D) Western blot analysis of PCBP1 in Smad3 immunoprecipitates from HeLa cells using conditions identical to panel (C). (E) Representative immunofluorescence staining of Smad3 (green) and PCBP1 (red) in cultured MSNs exposed to consecutive stimulation (same treatment in both treatment days) with activin A (Act), SKF81297 (SKF) or their combination as indicated. Counterstaining with MAP2 and DAPI is shown in the lower row. Scale bar, 25μM. (F, G) Quantification of PCBP1^+^ (F) or Smad3^+^ (G) nuclei in cultured MSNs after the indicated treatments (same treatment in both treatment days). Results are expressed as average ± SEM of % of MAP2^+^ cells displaying PCBP1 (F) or Smad3 (G) in the nucleus. ***, p<0.001; N=5; one-way ANOVA followed with Tukey’s multiple comparisons test. (H) Quantification of PCBP1/Smad3 co-localization in nuclei of cultured MSNs using conditions identical to panel (F). Results are expressed as average ± SEM of % of MAP2^+^ cells displaying co-localization of PCBP1 and Smad3 in the nucleus. (I) RT-qPCR analysis of PCBP1 mRNA in cultured MSNs exposed to consecutive stimulation as indicated (same treatment in both treatment days). Results are expressed as average ± SEM of fold change over vehicle. No significant differences were found by one-way ANOVA followed with Tukey’s multiple comparisons test (p≥0.05; N=4). (J) Representative images of PLA (red) between Smad3 and PCBP1 (S3/PCBP1) in cultured MSNs exposed to consecutive stimulation (same treatment in both treatment days) with activin A (Act), SKF81297 (SKF), PKA inhibitor (PKI) or SB431542 (SB) as indicated. Counterstaining with MAP2 and DAPI is shown in the lower row. Scale bar, 25μM. (K) Quantification of PLA between Smad3 and PCBP1 in cultured MSNs after the indicated treatments. Results are expressed as average ± SEM of number of PLA dots per cell. *, p<0.05, **, p<0.01; ***, p<0.001; N=5; one-way ANOVA followed with Tukey’s multiple comparisons test.

Importantly, two consecutive co-stimulations with SKF81297 and activin A induced a strong co-localization of PCBP1 and Smad3 in MSN nuclei, while individual treatment with either agent had a much lower effect (Fig. 2E, H). We note that the level of PCBP1 mRNA in MSNs was not affected by any of these treatments (Fig. 2I). To assess direct PCBP1/Smad3 interaction in neurons, we used the Proximity Ligation Assay (PLA) in primary cultures of MSNs treated with SKF81297 and activin A. Two consecutive co-stimulations induced a pronounced increase in PLA signals in MSNs, with little effect of either agent on its own (Fig. 2J, K). Treatment with the SB431542 or PKI14-22 inhibitors completely abolished the synergistic effect of SKF81297 and activin A on PCBP1/Smad3 interaction (Fig. 2J, K), suggesting that concurrent phosphorylation of PCBP1 and Smad3 by PKA and ALK4 kinases, respectively, is required for their interaction in MSN nuclei.

### PCBP1 and Smad3 interact with regulated intron 4 and adjacent sequences in FosB mRNA upon coincident activation of D1 and ALK4 receptors in NAc MSNs

RNA splicing factors may interact directly or indirectly with their target pre-mRNAs, but do not typically remain bound to the mature mRNA after splicing. In order to determine whether PCBP1 and Smad3 interact with FosB and ΔFosB mRNAs, we assessed the presence of these transcripts by quantitative RT-PCR in PCBP1 and Smad3 immunoprecipitates from extracts of HeLa cells after two consecutive stimulations with activin A, forskolin or both. No ΔFosB mRNA could be detected under any condition in either PCBP1 or Smad3 immunoprecipitates (N=3), suggesting that these proteins do not associate with the ΔFosB transcript. Very low levels of FosB mRNA could be detected in PCBP1 and Smad3 immunoprecipitates of untreated cells, while a moderate increase was observed after either forskolin or activin A treatment, respectively (Fig. 3A, B). On the other hand, two consecutive co-stimulations of HeLa cells with forskolin and activin A strongly potentiated the amount of FosB mRNA recovered from both PCBP1 and Smad3 immunoprecipitates (Fig. 3A, B), suggesting that the two pathways synergize to induce the association of these proteins with FosB mRNA. This question was addressed further using a modified PLA protocol developed to detect RNA/protein interactions (RNA-PLA) (Roussis et al., 2016; Zhang et al., 2016). We used antisense probes designed to recognize different regions in FosB mRNA, including a region in exon 4 (pE4) immediately upstream of intron 4 (i.e. the regulated intron), intron 4 (prI4), exon 3 (pE3) and a probe against the exon 4a/4b (pE4a/b) junction that forms after removal of intron 4, which is present in ΔFosB but not in FosB mRNA (Fig. 3C). Two consecutive co-stimulations of HeLa cells with forskolin and activin A produced robust interaction of PCBP1 and Smad3 with probes directed to exon 4 and intron 4 of FosB mRNA (Fig. 3D-F and S2A-C), but not with probes directed to exon 3 nor the exon 4a/4b junction (Fig. S3D, E). Stimulation with either forskolin or activin A on their own had little effect (Fig. 3D-F and S3A-C). The interactions of PCBP1 and Smad3 with FosB mRNA sequences were completely eliminated by treatment with PKA or ALK4 kinase inhibitors (Fig. 3D-F and S3A-C). No RNA-PLA signals were detected with sense probes for either exon 4 or intron 4 (Fig. S3D, E), nor with antisense probes and an unrelated antibody (Fig. S3F). Importantly, comparable results for both PCBP1 and Smad3 were obtained in primary cultures of NAc MSNs using the antisense probe against exon 4 (pE4) of FosB mRNA (Fig. 3G-I).

**Figure 3.**
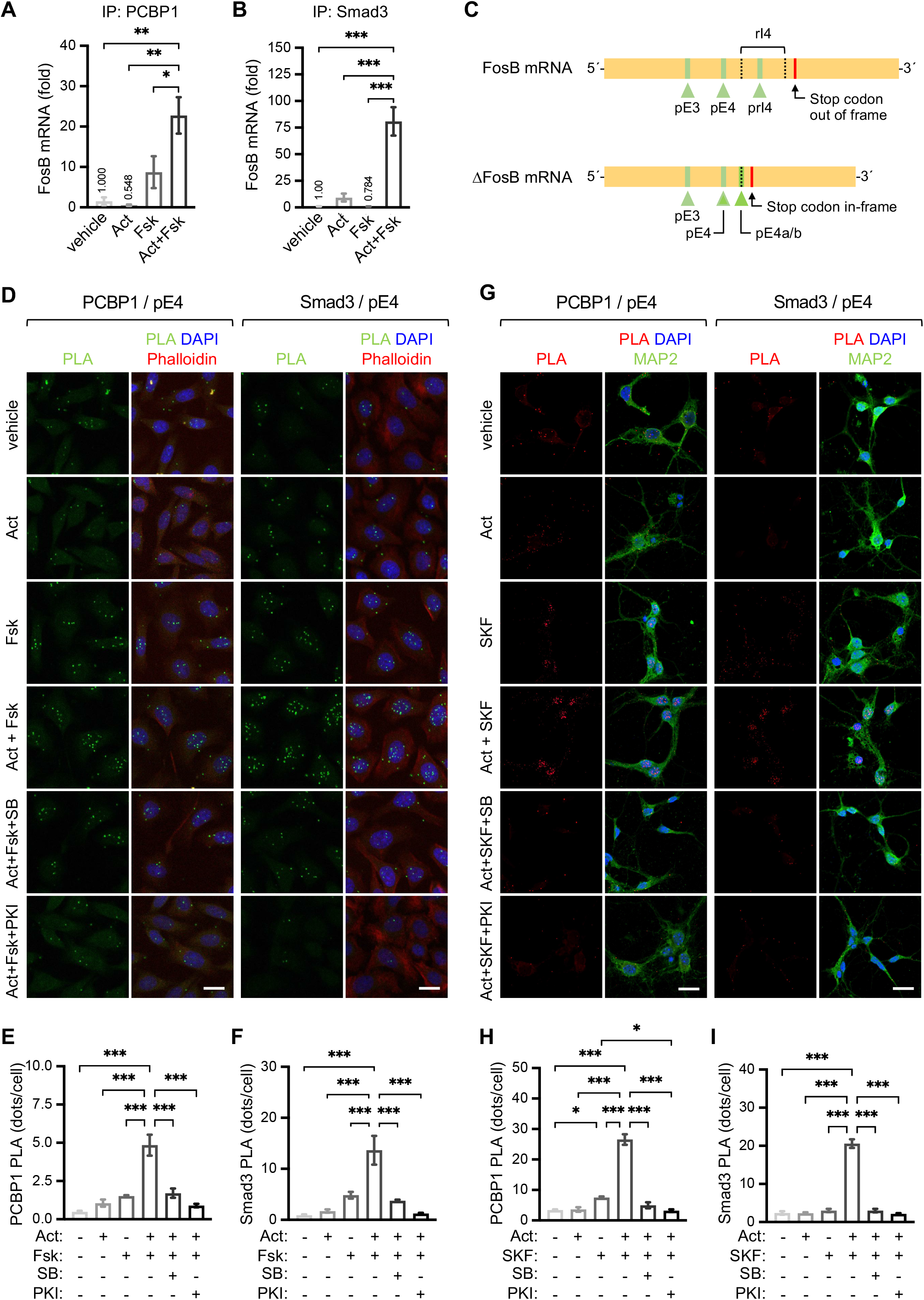
PCBP1 and Smad3 interact with regulated intron 4 and adjacent sequences in FosB mRNA upon coincident activation of D1 and ALK4 receptors in NAc MSNs. (A, B) RT-qPCR analysis of FosB mRNA in PCBP1 (A) and Smad3 (B) immunoprecipitates from HeLa cells exposed to consecutive stimulation (same treatment in both treatment days) with activin A (Act), forskolin (Fsk) or their combination as indicated. Results are expressed as average ± SEM of fold increase over vehicle . *, p<0.05; **, p<0.01; ***, p<0.001; N=4; one-way ANOVA followed with Tukey’s multiple comparisons test. (C) Schematic showing the location of probes used for RNA-PLA studies on the interaction of PCBP1 and Smad3 with sequences in FosB and ΔFosB mRNAs. The location of the regulated intron 4 (rI4) subjected to alternative splicing in FosB mRNA is indicted with black dotted lines. In ΔFosB mRNA, the dotted line indicates the junction between exons 4a and 4B after rI4 splicing. The locations of probes for exon 3 (pE3), exon 4 (pE4), regulated intron 4 (prI4) and exon 4a/4b junction (pE4a/b) are indicated in green. The red bar shows the location of the stop codon that becomes in-frame in ΔFosB mRNA after rI4 splicing. (D) Representative images of RNA-PLA (green) between PCBP1 (left) or Smad3 (right) and sequences in exon 4 (pE4) of FosB mRNA, respectively, in HeLa cells exposed to consecutive stimulation (same treatment in both treatment days) as indicated. Counterstaining with phalloidin and DAPI is shown in the second and fourth columns. Scale bars, 20μM. (E, F) Quantification of RNA-PLA between PCBP1 (E) or Smad3 (F) and pE4 in HeLa cells after the indicated treatments. Results are expressed as average ± SEM of number of RNA-PLA dots per cell. Only nuclear dots were quantified in this analysis. ***, p<0.001; N=4; one-way ANOVA followed with Tukey’s multiple comparisons test. (G) Representative images of RNA-PLA (green) between PCBP1 (left) or Smad3 (right) and sequences in exon 4 (pE4) of FosB mRNA, respectively, in cultured MSNs exposed to consecutive stimulation as indicated. Counterstaining with phalloidin and DAPI is shown in the second and fourth columns. Scale bars, 20μM. (H, I) Quantification of RNA-PLA between PCBP1 (E) or Smad3 (F) and pE4 in cultured MSNs after the indicated treatments. Results are expressed as average ± SEM of number of RNA-PLA dots per cell. Only nuclear dots were quantified in this analysis. *, p<0.05***; p<0.001; N=4; one-way ANOVA followed with Tukey’s multiple comparisons test.

### PCBP1 is required for the synergistic effects of dopamine and ALK4 signaling on ΔFosB mRNA induction and nuclear translocation of ΔFosB protein in NAc MSNs

In order to assess the requirement of PCBP1 for the induction of ΔFosB mRNA in response to synergistic activation of D1 and ALK4 receptors, we developed a lentiviral vector expressing shRNA targeting PCBP1 mRNA. In primary cultures of MSNs neurons, infection with shPCBP1 lentivirus reduced PCBP1 mRNA expression by 90%, while a lentivirus carrying a scrambled shRNA had no effect (Fig. 4A). In agreement with our previous findings, two consecutive co-stimulations of NAc MSNs with SKF81297 and activin A produced a robust induction of ΔFosB mRNA (>17-fold vs. vehicle), while SKF81297 on its own had a much lower effect (3-fold) (Fig. 4B). Interestingly, knock-down of PCBP1 strongly blunted the response of MSNs to consecutive co-stimulation with SKF81297 and activin A, but had no effect on the response to SKF81297 in the absence of activin A (Fig. 4B). In contrast, FosB mRNA levels were unaffected by PCBP1 knock-down regardless of treatment (Fig. 4C). Knock-down of PCBP1 also eliminated the synergistic effects of SKF81297 and activin A on the nuclear translocation of ΔFosB protein in MSNs (Fig. 4D, E), phenocopying the response of neurons lacking ALK4 (Fig. 1L). The effect of SKF81297 in the absence of activin A was again unaffected by PCBP1 know-down (Fig. 4E).

**Figure 4.**
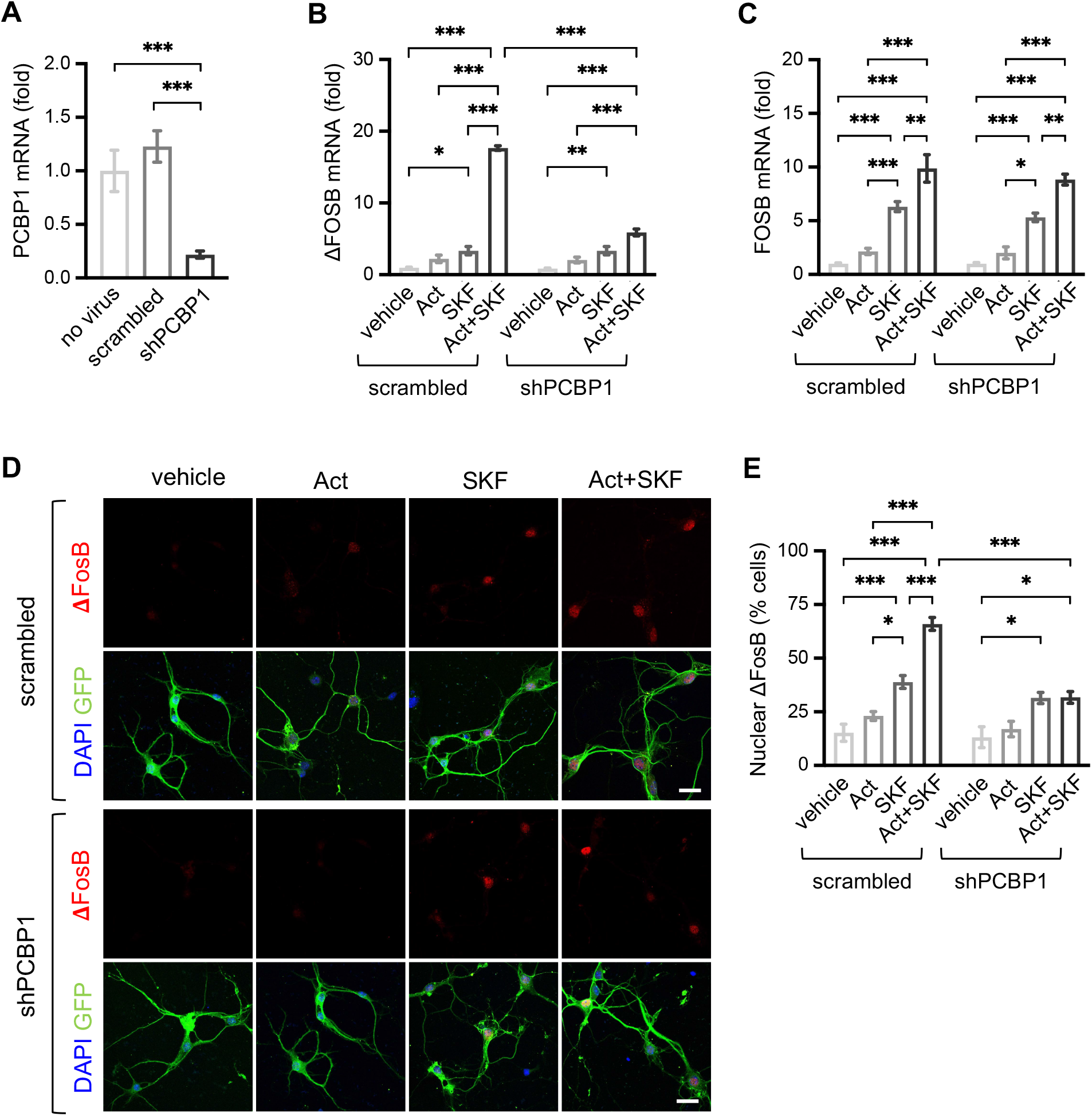
PCBP1 is required for the synergistic effects of dopamine and ALK4 signaling on ΔFosB mRNA induction and nuclear translocation of ΔFosB protein in NAc MSNs. (A) RT-qPCR analysis of PCBP1 mRNA in cultures of naive MSNs (no virus) or MSNs treated with lentiviruses expressing shRNA against PCBP1 (shPCBP1) or a scrambled shRNA. Results are expressed as average ± SEM of fold increase over control uninfected cells . ***, p<0.001; N=4; one-way ANOVA followed with Tukey’s multiple comparisons test. (B, C) RT-qPCR analysis of ΔFosB (B) and FosB (C) mRNA in cultured MSNs treated with scrambled and shPCBP1 lentiviruses and exposed to consecutive stimulation (same treatment in both treatment days) with activin A (Act), SKF81297 (SKF) or their combination as indicated. Results are expressed as average ± SEM of fold increase over vehicle . *, p<0.05; **, p<0.01; ***, p<0.001; N=4; two-way ANOVA followed with Tukey’s multiple comparisons test. (D) Representative immunofluorescence staining of ΔFosB (red) in cultured MSNs treated with scrambled and shPCBP1 lentiviruses and exposed to consecutive stimulation (same treatment in both treatment days) with activin A (Act), SKF81297 (SKF) or their combination as indicated. Counterstaining with green fluorescent protein (GFP, expressed by the lentiviruses) and DAPI is shown in the second and fourth rows. Scale bar, 25μM. (E) Quantification of ΔFosB^+^ nuclei in cultured MSNs treated with scrambled and shPCBP1 lentiviruses followed by the indicated treatments. Results are expressed as average ± SEM of % of GFP^+^ cells displaying ΔFosB in the nucleus. *, p<0.05; ***, p<0.001; N=4; two-way ANOVA followed with Tukey’s multiple comparisons test.

### ALK4 is required in NAc MSNs of adult mice for behavioral sensitization to cocaine

The above results show that ALK4 and dopamine signaling synergize in an *in vitro* paradigm of sensitization based on primary cultures of NAc MSNs. This effect was manifested by a greatly potentiated induction of ΔFosB mRNA, as well as nuclear translocation of ΔFosB protein, following two consecutive co-stimulations of ALK4 and D1 receptors. Based on these results, we investigated the importance of ALK4 signaling in an *in vivo* paradigm of sensitization to cocaine known as the “Two-Injection Protocol of Sensitization” or TIPS (Fig. 5A). TIPS has been widely used to study behavioral sensitization to a variety of addictive stimuli, including psychoactive drugs, such as amphetamine, morphine, nicotine and cocaine (Bernardi and Spanagel, 2014; Valjent et al., 2010). Locomotor behavior in response to cocaine injection was first studied in *Gad67*^Cre^;*Alk4*^fl/fl^ mutant mice. A single injection of cocaine (sensitization) induced a robust locomotor response of similar magnitude in both mutant and control mice compared to animals that received saline, indicating that ALK4 is not required for acute locomotor responses to this drug (Fig. 5B, left). Seven days after the first injection, saline and cocaine groups were divided into two subgroups, each of which received a second injection of either saline or cocaine (expression). Consecutive cocaine injections induced a strong potentiation of the locomotor response in control mice, doubling the distance travelled by the mice that received cocaine only the second time (Fig. 5B, right), indicative of successful sensitization. In contrast, ALK4 mutant mice showed no potentiation following consecutive cocaine administrations (Fig. 5B); i.e. their locomotor response to the second injection was the same regardless whether they received saline or cocaine in the first injection. These data indicate that induction of behavioral sensitization to cocaine requires a functional ALK4 receptor in GABAergic neurons. As a control experiment, targeting ALK4 recombination in GABAergic neurons derived from the medial ganglionic eminence (using *Nkx2.1*^Cre^;*Alk4*^fl/fl^ mutant mice), which comprise the majority of interneurons in cortex and hippocampus but not NAc MSNs (Xu et al., 2008), had no effect on Acvr1b mRNA expression in the NAc (Fig. S4A), nor did it affect cocaine sensitization and locomotor responses in the TIPS paradigm (Fig. 5C). In order to more specifically assess the importance of ALK4 in NAc MSNs, we used a *Gpr101*^Cre^ transgenic line that specifically targets Cre recombinase to these neurons (Reinius et al., 2015). These mutant mice (*Gpr101*^Cre^;*Alk4*^fl/fl^) showed a pronounced reduction in Acvr1b mRNA expression in NAc and striatum, but not in primary motor cortex (Fig. S4B-D). Similar to *Gad67*^Cre^;*Alk4*^fl/fl^ mice, *Gpr101*^Cre^;*Alk4*^fl/fl^ mutant mice lacking ALK4 specifically in NAc MSNs showed no cocaine sensitization; i.e. their locomotor response to a second cocaine injection was the same irrespectively of the nature of the first injection (Fig. 5D), strongly arguing for a specific requirement of ALK4 in NAc MSNs for behavioral sensitization to cocaine. In order to evaluate the acute requirement of ALK4 in sensitization to cocaine independently of possible developmental effects, we stereotaxically delivered adeno-associated viruses (AAV2) expressing Cre and GFP, or GFP alone, to the NAc of adult *Alk4*^fl/fl^ mice (Fig. S4E). The CRE-GFP virus reduced expression of Acvr1b mRNA by 60% in the NAc of these mice compared to control virus expressing only GFP (Fig. S4F) but did not affect Acvr1b mRNA expression in motor cortex (Fig. S4G). Importantly, injection of CRE-GFP virus in the NAc of *Alk4*^fl/fl^ mice abolished locomotor potentiation induced by consecutive cocaine administration in the TIPS paradigm, although it had no effect on the acute effects to a single injection (Fig. 5E). Control GFP virus did not affect locomotion (Fig. 5E). This result demonstrates that ALK4 is required at the time of drug exposure for behavioral sensitization to cocaine.

**Figure 5.**
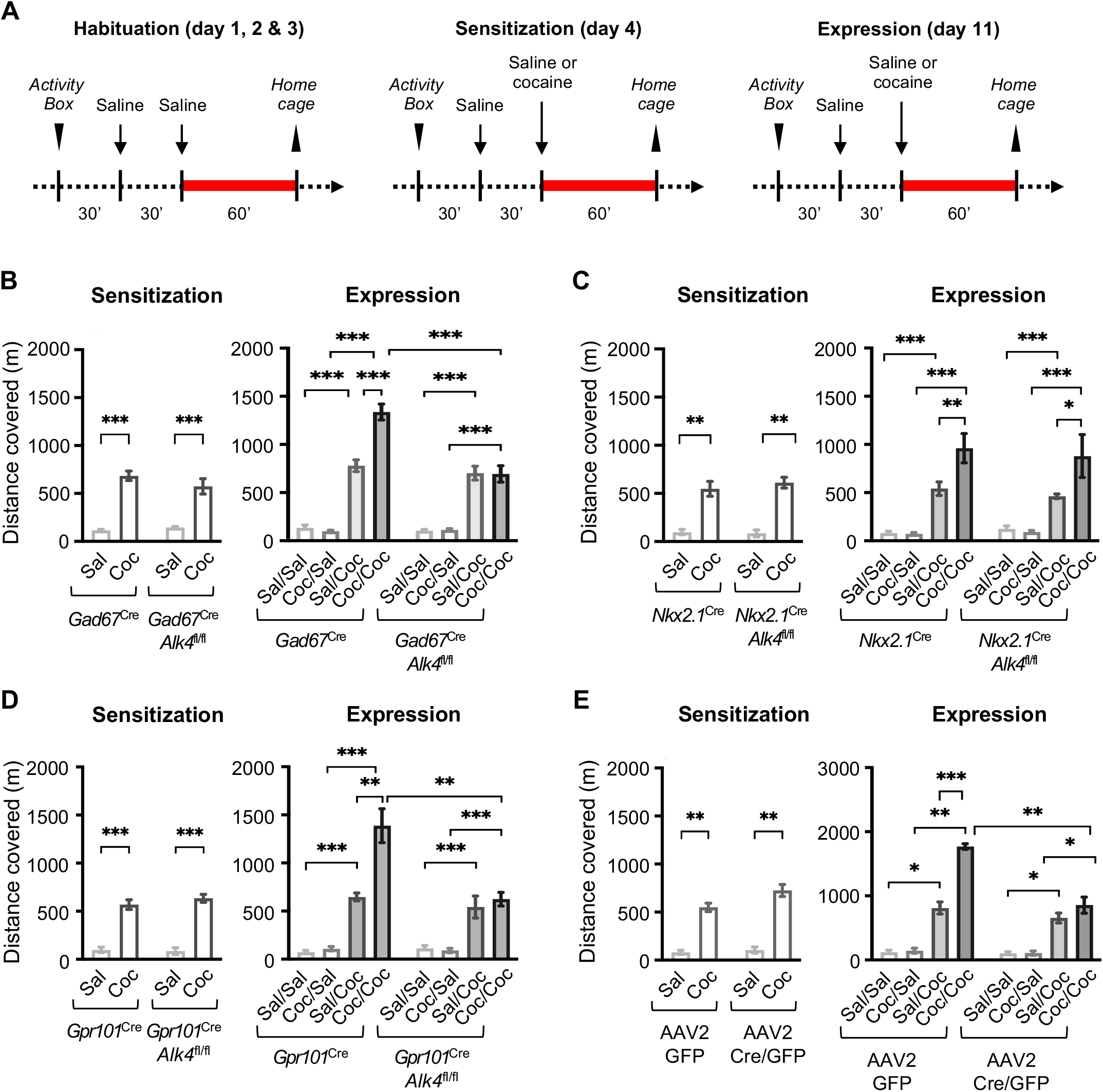
ALK4 is required in NAc MSNs of adult mice for behavioral sensitization to cocaine. (A) Schematic of the Two Injection Protocol of Sensitization (TIPS) for the habituation, sensitization and expression sessions. After entering the activity box, animals remain for 30 minutes before an intraperitoneal injection of saline. Thirty minutes later, saline (Sal) or cocaine (Coc) are injected after which locomotor activity is recorded during 60 minutes (red bars). Animals are returned to their home cage at the end of this period. This protocol generates 4 groups of animals, namely Sal/Sal, Sal/Coc, Coc/Sal and Coc/Coc according to what they received during the 1^st^ and 2^nd^ injections, respectively. (B) Locomotor responses of control mice (*Gad67*^Cre^) and mutant mice lacking ALK4 in GABAergic neurons (*Gad67*^Cre^;*Alk4*^fl/fl^) during cocaine TIPS. “Sensitization” denotes locomotor activity recorded after a first injection of saline (Sal) or cocaine (Coc), while “Expression” denotes locomotor activity in the 4 groups of mice after the second injection. Results are expressed as average ± SEM of distance covered in meters during the 60 minute recording period. ***, p<0.001; N=6; two-way ANOVA followed with Tukey’s multiple comparisons test. (C) Locomotor responses of control mice (*Nkx2.1*^Cre^) and mutant mice lacking ALK4 in GABAergic neurons derived from the medial ganglionic eminence (*Nkx2.1*^Cre^;*Alk4*^fl/fl^) during cocaine TIPS using conditions identical to panel (B). *, p<0.05; **, p<0.01; ***, p<0.001; N=6. (D) Locomotor responses of control mice (*Gpr101*^Cre^) and mutant mice lacking ALK4 in NAc MSNs (*Gpr101*^Cre^;*Alk4*^fl/fl^) during cocaine TIPS using conditions identical to panel (B). **, p<0.01; ***, p<0.001; N=5. (E) Locomotor responses of *Alk4*^fl/fl^ mice stereotaxically injected in NAc with control AAV2 viruses expressing only GFP (AAV2 GFP) or AAV2 viruses expressing both Cre recombinase and GFP (AAV2 Cre/GFP) during cocaine TIPS using conditions identical to panel (B). *, p<0.05; **, p<0.01; ***, p<0.001; N=6.

### Activin A is induced in NAc microglia after cocaine TIPS

We assessed the expression of endogenous ALK4 ligands in the NAc and their possible regulation upon cocaine sensitization. We found expression of Inhba mRNA (encoding activin A) and Gdf-1 mRNA (encoding GDF-1) in several brain structures, including NAc/striatum (Fig. S5A, B). In the NAc, activin A immunoreactivity was detected among DARPP32^+^ neurons, GFAP^+^ astrocytes and Iba1^+^ microglia cells (Fig. S5C). Strong immunoreactivity for activin A was found in microglia cell bodies of both NAc and striatum (Fig. S5D). A single injection of cocaine produced a moderate increase in NAc Inhba mRNA which did not reach statistical significance (Fig. S5E). However, Inhba mRNA was significantly upregulated in the NAc of mice that received two consecutive cocaine injections according to the TIPS protocol (Fig. S5F), indicating that cocaine sensitization, but not a single acute exposure, is able to increase Inhba mRNA in the NAc. At the protein level, the number of cells expressing activin A increased significantly among the Iba1^+^ population (Fig. S5G, H), but not among DARPP32^+^ neurons (Fig. S5I, J) in the mice that received two consecutive cocaine injections. These results suggest that NAc microglia represent an important source of activin A ligand upon sensitization to cocaine.

### ALK4 is required in NAc MSNs of adult mice for sustained induction of ΔFosB mRNA during cocaine sensitization

As expected, acute induction of immediate early genes FosB and c-Fos could be readily detected in NAc of both control and *Gad67*^Cre^;*Alk4*^fl/fl^ and *Gpr101*^Cre^;*Alk4*^fl/fl^ mutant mice 1h after the second cocaine injection (Fig. 6A, B, D, E). However, the levels of ΔFosB mRNA remained unchanged at this time point regardless of treatment or genotype (Fig. 6C, F). Expression of FosB and c-Fos mRNAs returned to basal levels 24h after the second injection in the NAc of both control and mutant mice (Fig. 6G, H, J, K). In contrast, a strong induction of ΔFosB mRNA was observed 24h after the second injection in the NAc of control mice that received two consecutive cocaine doses (Fig. 6I, L). Importantly, this response was completely abolished in both lines of mutant mice (Fig. 6I, L). Similar to ΔFosB, the levels of ARC and EGR1 mRNAs were also greatly induced 24h after the second cocaine injection (Fig. S6A-D). However, mutant mice showed no lasting induction of ARC and EGR1 mRNAs 24h after cocaine TIPS (Fig. S6A-D). Together, these results indicate a profound deficit in long-lasting induction of ΔFosB and its downstream targets ARC and EGR1 in NAc after specific ablation of ALK4 in MSNs.

**Figure 6.**
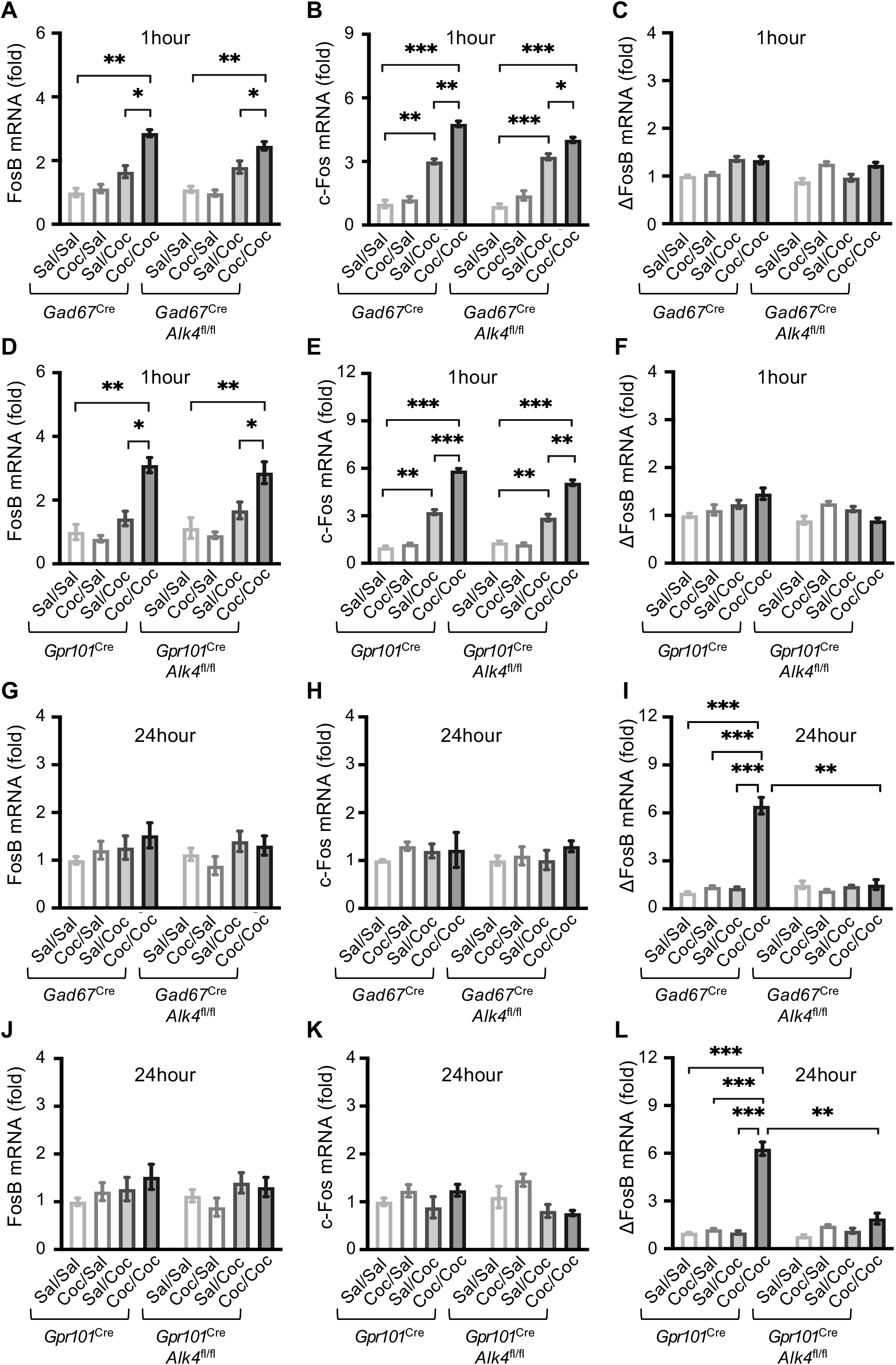
ALK4 is required in NAc MSNs of adult mice for sustained induction of ΔFosB mRNA during cocaine sensitization. (A-F) RT-qPCR analysis of FosB (A, D), c-Fos (B, E) and ΔFosB (C, F) mRNAs in NAc of *Gad67*^Cre^;*Alk4*^fl/fl^ (A-C) and *Gpr101*^Cre^;*Alk4*^fl/fl^ (D-F) mutant mice 1 hour after the second saline (Sal) or cocaine (Coc) injection of TIPS. Results are expressed as average ± SEM of fold increase over the saline group (Sal/Sal) of control mice (*Gad67*^Cre^ and *Gpr101*^Cre^, respectively). *, p<0.05; **, p<0.01; ***, p<0.001; N=5; two-way ANOVA followed with Tukey’s multiple comparisons test. (G-L) RT-qPCR analysis of FosB (G, J), c-Fos (H, K) and ΔFosB (I, L) mRNAs in NAc of *Gad67*^Cre^;*Alk4*^fl/fl^ (G-I) and *Gpr101*^Cre^;*Alk4*^fl/fl^ (J-L) mutant mice 24 hour after the second saline (Sal) or cocaine (Coc) injection of TIPS using conditions identical to panels A-F, respectively. **, p<0.01; ***, p<0.001; N=5; two-way ANOVA followed with Tukey’s multiple comparisons test.

## Discussion

Since its discovery two decades ago (Mumberg et al., 1991; Nakabeppu and Nathans, 1991), ΔFosB has been the subject of numerous investigations, with hundreds of papers describing various aspects of its induction, expression and downstream effects. The relationship between Fos family proteins and activation of D1 receptors is equally old (Hope et al., 1992; Young et al., 1991); and the association of ΔFosB with chronic cocaine administration was also uncovered at about the same time (Hope et al., 1994). Despite this large body of research, the molecular mechanisms that promote the alternative splicing of FosB mRNA to generate ΔFosB have been unknown. The current view is that ΔFosB mRNA is generated constitutively above a certain level of FosB mRNA expression (Alibhai et al., 2007). According to this model, the machinery that promotes the alternative splicing of FosB mRNA is not thought to be affected or regulated by dopamine signaling. In the present study, we describe a previously unknown mechanism by which dopamine signaling synergizes with the TGFβ superfamily receptor ALK4 to potentiate the production of ΔFosB in MSNs of the NAc through activation of the RNA-binding protein PCBP1. In the absence of ALK4, locomotor responses to a second dose of cocaine were identical to the first, and long lasting ΔFosB mRNA expression was abolished, indicating that persistent expression of ΔFosB mRNA and behavioral sensitization to cocaine are both dependent on ALK4 signaling in NAc MSNs of adult mice. Given the critical role of ΔFosB in reward responses and addiction, these findings establish a new paradigm for understanding the early molecular events leading to the structural and functional adaptations that underlie those processes.

The molecular mechanisms that mediate the alternative splicing of FosB mRNA to generate ΔFosB are understudied and remain elusive. PCBP1 (poly-(rC)-binding protein 1, also known as hnRNPE1) has been known to participate in various mRNA processing steps, including alternative splicing, stability and translational repression (Hwang et al., 2017; Meng et al., 2007). Phosphorylation of PCBP1 at Thr^60^ and Thr^127^ by Pak1 was shown to induce its translocation to the nucleus, where it regulated the alternative splicing of CD44 mRNA, promoting inclusion of its variable exon (Meng et al., 2007). Subsequent studies found that PCBP1 can be phosphorylated at these and other sites by multiple Ser-Thr kinases, including AKT, CDKs and MAPKs, following activation of growth factor receptors (Chaudhury et al., 2010; Tripathi et al., 2016). In conjunction with TGFβ signaling, activated PCBP1 and Smad3 were shown to form a complex that translocated to the cell nucleus where it promoted exclusion of CD44 mRNA regulated exon, contributing to tumor formation (Tripathi et al., 2016). In the present studies, we found that PCBP1 and Smad3 can be activated by PKA and ALK4, respectively, inducing their association and nuclear translocation in NAc MSNs, thereby extending this paradigm to D1 and ALK4 receptor signaling in neuronal cells. The ability of activin A to potentiate PCBP1 nuclear translocation induced by D1 receptor activation, suggests that activated Smad3 and PCBP1 first interact in the cytoplasm, from where Smad3 nuclear localization signals may then facilitate the nuclear translocation of the complex.

We found that PCBP1 and Smad3 interacted with FosB mRNA at and around intron 4 sequences, but not other regions in FosB mRNA, nor the exon 4a/4b junction present in ΔFosB mRNA but absent in FosB, suggesting the involvement of these proteins in FosB intron 4 splicing rather than other processes, such ΔFosB mRNA stabilization. We note that, although activin A greatly potentiated the effects of SKF81297 on ΔFosB mRNA induction and ΔFosB nuclear translocation, SKF81297 was also able to enhance both these processes on its own, albeit at more moderate levels. In contrast to the co-stimulation regime, however, SKF81297 induced FosB and ΔFosB expression to comparable degrees when administered in the absence of activin A, in agreement with the prevalent model of constitutive ΔFosB generation proportionally to FosB mRNA (Alibhai et al., 2007; Nestler, 2008; 2013). On the other hand, the fact that PCBP1 knock-down in MSNs specifically interfered with the potentiated induction of ΔFosB mRNA afforded by co-stimulation with activin A and SKF81297 without affecting FosB mRNA levels, suggests a direct role for PCBP1 in enhancing ΔFosB mRNA generation independently of FosB gene transcription. In agreement with this, PCBP1 knock-down did not impact the effects of SKF81297 on ΔFosB expression and nuclear localization in the absence of activin A, indicating that PCBP1 specifically participates in the synergy between D1 with ALK4 signaling that enhances ΔFosB mRNA levels, rather than the basal mechanism of ΔFosB mRNA generation. Similar to its role in the alternative splicing of CD44 mRNA (Meng et al., 2007; Tripathi et al., 2016), the PCBP1/Smad3 complex may regulate the function of components of the basal splicing machinery, such as factors associated with the U2 small nuclear ribonucleoprotein complex, to enhance splicing of FosB mRNA intron 4, thereby increasing the generation of ΔFosB mRNA.

Psychostimulants of abuse, including cocaine, methamphetamine and ecstasy, have been shown to induce neuroinflammation in the brain through activation of resident endothelial, glial and immune cells (Clark:2013dg; Anderson and Döffinger, 2017). In vitro and in vivo studies have provided evidence indicating that repeated exposure to cocaine can induce activation of microglial cells, particularly in striatum, through a mechanism involving increased production of reactive oxygen species (ROS) and inflammasome activity (Burkovetskaya et al., 2020; Chivero et al., 2021; Liao et al., 2016). In our studies, we found significantly increased activin A production in NAc microglia following two consecutive cocaine exposures, but not after a single administration. An earlier study reported elevated levels of activin A in the NAc of rats after a cocaine relapse (Wang et al., 2017). Increased activin A production in NAc microglia after repeated cocaine exposure suggests its upregulation may be a consequence of cocaine-mediated neuroinflammation.

In conclusion, we propose that the synergistic effects of D1 and ALK4 signaling mediated by concurrent activation of PCBP1 and Smad3 in NAc MSNs represent a previously unrecognized coincidence-detection mechanism that potentiates ΔFosB production and reward-related behavior in response to repeated exposure to particularly salient stimuli. We postulate that such mechanism facilitates sensitization to specific stimuli only when these are of sufficient magnitude, duration and persistence to activate concurrent D1 and ALK4 signaling in NAc MSNs. Targeting the key components of this mechanism may lead to interventions that counteract the effects of drugs of abuse.

## Methods

### Mice

Mice were housed 2-5 per cage in standard conditions in a temperature and humidity-controlled environment, on a 12/12 h light/dark cycle with ad libitum access to food and water. The following transgenic mouse lines were used for experiments: *Gad67*^Cre^ (Tolu et al., 2010), *Nkx2.1*^Cre^ (Xu et al., 2008), *Gpr101*^Cre^ (Reinius et al., 2015) and *Alk4*^fl/fl^ (Göngrich et al., 2020). All animals were bred in a C57BL6/J background (Jackson). Both males and females were used in the studies; mice were 12 to 14 week old at the start of the experiments. Animal protocols were approved by Stockholms Norra Djurförsöksetiska Nämnd and are in accordance with the ethical guidelines of Karolinska Institutet (ethical permits N173-15; 11563-2018 and 11571-2018 and N68/16 following EU directive 2010/63/EU).

### Cell culture and treatments

Medium Spiny Neuron (MSN) culture was performed according to Penrod et al., 2011 (Penrod et al., 2011) with minor modifications. Briefly, pregnant dames were killed by cervical dislocation at 16.5 days of gestation, embryos were collected and brains were removed in ice-cooled Ca^+2^/Mg^+2^-free Hank’s balanced salt solution (Gibco) with 10mM HEPES (Thermofisher). After removal of all meninges, ganglionic eminences, i.e protostriatum, were dissected in fresh Hank’s/HEPES solution. Collected tissue was transferred to 15ml tubes and digested with 0.25% Trypsin-EDTA (Gibco) during 30 min at 37°C. Trypsin was inactivated with 25% Newborn Calf Serum (NCS, Gibco) and cells were dissociated by 10 passages through a glass pipette in the presence of 25U DNase (Roche). Cells were then collected by centrifugation for 5 min at 500xg, resuspended in plating medium: MEM containing Earĺs salts (Gibco), supplemented with 10mM HEPES, 10mM sodium pyruvate, 0.5mM glutamine, 12.5μM glutamate, 0.6% glucose and 10% NCS. Cells were centrifuged one more time, and again resuspended in plating medium. Cells were then plated at a density of 30,000 cells per coverslip in 24-well plates; coverslips were coated with 100 µg/ml poly-D-lysine (Sigma-Aldrich) and 4µg/ml mouse Laminin (Cultrex, RnD). One to 3 hours later, after cells had adhered to the coverslips, plating medium was replaced by conditioned growth medium: Neurobasal (Gibco), containing B27 supplement (Gibco) and 0.5 mM glutamine (Gibco) which that had been conditioned on a monolayer of glial cells (see below). One half of the medium was replaced after 3 days with fresh conditioned growth media. Cells were maintained for 7 days at 37°C in 95% O_2_/5% CO_2_ atmosphere before the start of the experiments.

For preparation of glial cell conditioned medium, cortices of postnatal day 1-4 (P1-4) mice with the striatal tissue removed. Minced tissue was incubated in Hank’s balanced salt solution supplemented with 0.25% Trypsin-EDTA and DNAse at 37°C for 30 minutes. After digestion, an equal volume of glia plating medium (MEM with 10mM HEPES, 1mM Sodium Pyruvate, 0.2mM glutamine, 10% NCS, 0.6% Glucose and 1X Penicillin-Streptomycin) was added to inhibit the trypsin and the tissue was collected by centrifugation at 500xg for 5 minutes. The tissue was resuspended in glia plating medium, mechanically dissociated using a flame-polished glass pipette and filtered through a 0.7μm cell strainer (Fisher scientific). Cells from one pup were plated onto 10 cm standard tissue culture dishes. Media was replaced one day after plating and once per week for each subsequent week. Conditioned medium was harvested after cell monolayers reached 70% confluence.

HeLa cells were grown in MEM supplemented with 2 mM glutamine (Gibco), 20 μg/ml penicillin/streptomycin (Gibco), and 10% heat inactivated Fetal bovine serum (FBS, Gibco). Cells were maintained at 37°C in a 95% O_2_/5% CO_2_ atmosphere. Cell line experiments were performed after cell monolayers reached 80% confluence.

MSN cultures (after 7 days in vitro) or cell lines (after 80% confluence) were treated for 2 hours with one or more of substances, as indicated in the Results section, at the following concentrations: Activin A (RnD) 100ng/ml, SKF91297 (Tocris) 1µM, Forskolin (Tocris) 25μM, SB41542 (Sigma-Aldrich) 10μM, PKI14,22 amide (Tocris) 1μM. For the sequential stimulation protocol, cells were washed after 2 hour treatment in phosphate buffer saline (PBS, Gibco) and fresh growth media was added. Two days later, cells were treated again for 2 hours and washed before fixation or lysis.

### RNA extraction and quantitative real-time quantitative PCR (RT-qPCR)

Total mRNA was isolated from cell monolayers and dissected brain areas using RNeasy Mini Kit (Qiagen) according to the manufacturer’s protocol. QIAshredder (Qiagen) was used for tissue homogenization prior RNA extraction. RNA purity and quantity of RNA were measured in a Nanodrop 1000 (ThermoScientific). cDNA was synthesized by reverse transcription using 150ng of RNA in a reaction volume of 20μl using High Capacity cDNA Reverse Transcription Kit (Applied Biosystems) according to the manufacturer’s protocol. cDNA samples were diluted 1:3 in distilled water before use. For measurement of 18S RNA levels, a further 1:800 dilution was made. The geometric mean of mRNAs encoding the ribosomal protein 18S was used for normalization. RT-qPCR was performed on the 7500 Real-Time PCR System (Applied Biosystems) with SYBR Green fluorescent probes and the following conditions: 40 cycles of 95°C for 15sec, 60°C for 1min and 72°C for 30sec. Forward and reverse primers were used at 100 nM each. The primer pairs used were as follows: m18s (Fw: 5’-GCAATTATTCCCCATGAACG-3’; Rv: 5’-GGCCTCACTAAAACCATCCAA-3’); h18s (Fw: 5’-CTCAACACGGGAAACCTCAC-3’; Rv: 5’-CGCTCCACCAACTAAGAACG-3’); ΔFosB (Fw: 5’-AGGCAGAGCTGGAGTCGGAGAT -3’; Rv: 5’-GCCGAGGACTTGAACTTCACTCG - 3’); FosB (Fw: 5’-GTGAGAGATTTGCCAGGGTC-3’; Rv: 5’-AGAGAGAAGCCGTCAGGTTG -3’); c-Fos (Fw: 5’-GGAATTAACCTGGTGCTGGA- 3’; Rv: 5’-AGAGAGAAGCCGTCAGGTTG -3’); ARC (Fw: 5’-CTTGCCTCCTGTCCTGAGC -3’; Rv: 5’-AGAAAGCAGCAGCAAGATGG -3’); EGR1 (Fw: 5’-AATCCCCCTTCGTGACTACC -3’; Rv: 5’-GCAAGGATGGAGGGTTGG -3’); Acvr1b (Fw: 5’-CCAACTGGTGGCAGAGTTAT-3’; Rv: 5’-CTGGGACAGAGTCTTCTTGATG-3’); PCBP1 (Fw: 5’-ATAGATGATTGGAACCAGGTAAAGT-3’; Rv: 5’-GCTCCAAGGGAGCAAGCGAGCAGT-3’); Inhba (Fw: 5’-ATCATCACCTTTGCCGAGTC-3’; Rv: 5’-ACAGGTCACTGCCTTCCTTG-3’); Gdf-1 (Fw: 5’-TTCTGCCAGGGCACGTGCG-3’; Rv: 5’-GGAGCAGCTGCGTGCATGAG-3’). Each experiment was carried out on biological triplicates for each data point. Data analysis was performed using the 2−ΔCt method, and relative expression levels were calculated for each sample after normalization against 18S RNA levels.

### Co-Immunoprecipitation, nuclear fractionation, Western blotting and RNA-immunoprecipitation

HeLa cell lysates were prepared on ice using RIPA buffer (Thermofisher) containing protease and phosphatase inhibitors (Roche). Briefly, growth media was removed, cells were washed twice with PBS, then lifted in RIPA buffer using a cell scraper. The samples were centrifuged for 10 minutes at 5,000xg and the supernatant was collected. Immunoprecipitation from cell lysates was performed using Dynabeads Protein A (Thermofisher) following the manufacturer ’s protocol. Protein concentration was measured with the Pierce™ BCA Protein Assay Kit (ThermoFisher), and 200μg of total protein lysate were used for each immunoprecipitation reaction. Rabbit anti PCBP1 (1:150, #85341, Cell signaling) and rabbit anti Smad3 (1:100, #9624, Cell signaling) were used for immunoprecipitation. Rabbit anti c-Jun (1:150, #9165, Cell Signaling) was used as mock control antibody. For Western blotting, immunoprecipitates were boiled at 95°C for 10 minutes in sample buffer (Life Technologies) before separation by SDS-PAGE in 4-15% Mini-PROTEAN® TGX™ Precast Gel (Bio-Rad). Proteins were transferred to polyvinylidene fluoride (PVDF) membranes (Amersham). Membranes were blocked with 5% non-fat milk in TBST and incubated with antibodies overnight in 1% milk. The antibodies used were as follows: mouse anti PCBP1 (1:400, sc-137249 Santa Cruz biotech), rabbit anti Smad3 (1:1000, #9513, Cell Signaling) and rabbits anti-phospho-PKA substrates (RRXS*/T*) (1:1000, #9624; Cell Signaling).

Cytosolic and nuclear protein fractions were prepared using NE-PER™ Nuclear and Cytoplasmic Extraction Reagents (ThermoFisher) following the manufacture’s protocol. Western blotting was performed as indicated above using 20μg of protein per sample. The antibodies used on membranes containing cytosolic and nuclear protein fractions were: rabbit anti PCBP1 (1:1000, #8534, Cell Signaling), mouse anti PCNA (1:1000, MAB424R, Sigma-Aldrich), αTubulin (1:2000, 05-829, Sigma-Aldrich). Immunoreactivity was detected using appropriate horseradish peroxidase (HRP)-conjugated secondary antibodies (1:5000, Dako). Immunoblots were developed using the Clarity Western ECL substrates (Bio-Rad) or the SuperSignal West Femto Maximum sensitivity substrate (ThermoFisher) and images were acquired with ImageQuant LAS4000 (GE Healthcare). Image analysis and quantification of band intensities were done with ImageQuant software (GE Healthcare).

RNA immunoprecipitation was performed using the Magna RIP® RNA-Binding Protein Immunoprecipitation Kit (Sigma-Aldrich) according to the manufacturer’s protocol. Approximately 2×10^7^ Hela cells were used for each immunoprecipitation. Mousse anti PCBP1(5μg, sc-137249 Santa Cruz Biotech) and anti-Smad3 (1:100, #9513, Cell Signaling) were used for immunoprecipitation as described above. Binding to ΔFosB or osSB mRNA was assessed by RT-qPCR as described above using the following primers: ΔFosB (Fw: 5’-AGGCAGAGCTGGAGTCGGAGAT -3’; Rv: 5’-GCCGAGGACTTGAACTTCACTCG -3’); FosB (Fw: 5’-GTGAGAGATTTGCCAGGGTC-3’; Rv: 5’-AGAGAGAAGCCGTCAGGTTG -3’).

### Immunocytochemistry and proximity ligation assay (PLA)

Coverslips containing cell monolayers were washed twice in PBS for 10 minutes and fixed for 15 minutes in 4%PFA (Histolab) / 4% Sucrose. Cells were washed 2 extra times to remove any remaining PFA. Cells were permeabilized in 0.3% Triton X-100 (Sigma-Aldrich) in PBS for 10 minutes, followed for 1 hour blocking in 5% Normal Donkey Serum (NDS, Jackson ImmunoResearch) and 0.05% Triton X-100 in PBS. Cells were then incubated overnight at 4°C under gentle agitation with the primary antibodies in 2% NDS, 0.05% Triton X-100 in PBS. Primary antibodies used were as follows: rabbit anti ΔFosB (1:500, #14695, Cell Signaling), mouse Anti-PCBP1 (1:400, sc-137249Santa Cruz biotech), rabbit anti Smad3 (1:250, #9513, Cell Signaling), chicken anti MAP2 (1:5000, ab5392, Abcam) or mouse anti MAP2 (1:200; M4403; Sigma-Aldrich). Coverslips were subsequently washed 3 times for 10 minutes and incubated with the appropriate secondary antibody for 2 hours at room temperature. Secondary antibodies were donkey anti-rabbit IgG Alexa Fluor 555, A31572; donkey anti-goat IgG Alexa Fluor 488, A11055; donkey anti-mouse IgG Alexa Fluor 488, A21202; donkey anti-mouse IgG Alexa Fluor 555, A31570; donkey anti-mouse IgG Alexa Fluor 647, A31571; donkey anti-goat IgG Alexa Fluor 555, A21432 (1:800, Life Technologies and Invitrogen); and donkey anti-chicken Alexa Fluor647 (1:750; 703-606-155, Jackson ImmunoResearch).

PLA was performed according to the manufacturer’s protocol (Sigma-Aldrich). Briefly, cell cultures were fixed as indicated above. Cells were washed 10 minutes in 0.2% Triton X-100 and blocked for 1 hour in 5% NDS and 0.05% Triton X-100 in PBS before incubation with rabbit anti Smad3 (1:250, #9624, Cell Signaling), mouse anti PCBP1 (1:200, sc-137249; Santa Cruz biotech) and chicken anti MAP2 (Abcam, ab5392 ,1:2000) overnight at 4°C with gentle agitation. Coverslips were subsequently washed with PBS, incubated with minus (anti-mouse) and plus (anti-rabbit) PLA probes for 1 hour at 37 °C, washed with buffer A, incubated with Ligation-Ligase solution for 30 minutes at 37°C, washed again with buffer A, and finally incubated with Amplification-Polymerase solution (Duolink™ In Situ detection reagents Orange) for 100 minutes at 37 °C. Alexa anti-chicken were included during the amplification step to reveal the MAP2 signal. After washing with buffer B, sections were incubated for 10 minutes with DAPI (1:10000 in buffer B) and mounted on glass slides with DAKO mounting media.

Our RNA-PLA protocol was adapted from Roussis et al., 2016 (Roussis et al., 2016) and Zhang et al., 2016 (Zhang et al., 2016). One of the primary antibody/probe pairs of standard PLA was replaced by a DNA oligonucleotide containing a 40–50 nucleotide (nt) segment complementary to the RNA of interest, 20–35 nt poly-A as a linker segment, and a common 25 nt segment that serves as one of the PLA hybridization/ligation arms (PLA PLUS probe). Briefly, cells were washed twice for 10 minutes in Dulbecco’s modified Phosphate Saline Buffer (DPBS, Thermofisher) and fixed with 4% PFA / 4% sucrose (Sigma-Aldrich) for 15 minutes. After two washes in DPBS, cells were permeabilized for 10 minutes with 0.2% Triton-X100, washed once in DPBS, and blocked in Blocking Buffer (10mM Magnesium-Acetate, 50mM Potassium acetate, 250mM NaCl, 2% BSA,0.05% Tween-20, 20 μg/ml Sheared Salmon Sperm DNA (Thermofisher) for 1 hour. Probes (100nM) were heated to 70°C for 5 minutes in blocking buffer and incubated with the samples overnight at 58°C with gentle agitation. The following probes were used: Exon 4: Antisense: 5’-GGCCCCTCTTCGTAGGGGATCTTGCAGCCCGGTTTGTGGGCCACC-3’; Sense: 5’-GGTGGCCCACAAACCGGGCTGCAAGATCCCCTACGAAGAGGGGCC-3’.

Regulated Intron 4: Antisense: 5’-CCGAAGCCGTCTTCCTTAGCGGATGTTGACCCTGGCA-3’; Sense: 5’-TGCCAGGGTCAACATCCGCTAAGGAAGACGGCTTCGG-3’. Exon 3: Antisense 5’-TCTCTGCGAACCCTTCGCTTTTCTTCTTCTTCTGGGGTA-3’. Exon 4a/b junction: Antisense: 5’-GGAAGGGGTCGCCGAGGACTTGAACCTCGGCCAGCGGGCCTGGCCC-3’. All sequences were followed at the 3’ with the PLA-chimera probe sequence: 5’-AAAAAAAAAAAAAAAAAAAAAAAAAAATATGACAGAACTAGACACTCTT-3’. After overnight incubation, cells were washed three times in DPBS and incubated in NDS containing 0.02% Triton-X100 for 1 hour. Primary antibodies were added in 2% NDS, 0.02% Triton-X100 and incubation proceeded at 4°C overnight with gentle agitation. Primary antibodies were used for RNA-PLA were: mouse Anti-PCBP1 (1:200, sc-137249 Santa Cruz biotech), rabbit anti Smad3 (1:250, #9513, Cell Signaling) and Sirt1 (1:250, #2028; Cell Signaling). Chicken anti MAP2 (1:5000, ab5392, Abcam) or mouse anti MAP2 (1:200; M4403; Sigma-Aldrich) were used for counterstaining of MSNs; Alexa Fluor A488 Phalloidin (1:1000, A12379, Thermofisher) or Alexa Fluor 568 (1:1000, A12380, Thermofisher) were used for counterstaining of HeLa cells. After overnight incubation, samples were washed twice with DPBS for 10 minutes. PLA reactions were then performed as described above.

### PCBP1 knock-down by lentivirus infection

Short interference mouse PCBP1 RNAi plasmids were purchased from Origene (TL502540). Lentiviral particles expressing the PCBP1 shRNAi or scramble control were produced by the VirusTech Core Facility, Department of Medical Biochemistry and Biophysics (MBB) Karolinska Institutet. Neurons were infected after 7 days in vitro with PCBP1-shRNAi or scrambled sc-shRNAi (MOI: 100) overnight, supplemented with 1 μg/ml polybrene (Sigma-Aldrich).

### Two-Injection Protocol of Sensitization (TIPS) to cocaine

TIPS was performed according to Valjent et al. (2010) (Valjent et al., 2010) with minor modifications. Briefly, mice were placed in the center of a 48cm x 48cm transparent acrylic box equipped with light-beam strips (TSE Actimot system) and allowed to move freely. Distance travelled was calculated using Actimot Software. All mice were habituated to the test apparatus, handling, and procedure for 3 consecutive days before the actual experiment. During the habituation procedure mice were placed for 30min in the activity box, received a first injection of saline (0,9% NaCl; Sal), were placed back in the box for 30 min, received a second saline injection, and were placed again for 1h in the box before returning to the home cage. For the first drug injection (day 4, “sensitization”), a similar handling was repeated, except that, the second saline injection was replaced by either a saline or 20mg/kg cocaine-HCl (Coc; Sigma-Aldrich), resulting in 2 different groups of mice, namely Sal and Coc, respectively. The mice were then placed back in the activity box for 1 hour to register their locomotor activity. Seven days later, Sal and Coc groups were further divided before receiving a second injection of saline or cocaine (day 11, “expression”), resulting in 4 groups of mice, namely Sal/Sal, Sal/Coc, Coc/Sal and Coc/Coc, respectively. The mice were then placed back in the activity box for 1 hour to register their locomotor activity and sacrificed after either 1 or 24 hours for immunohistochemistry or RNA analysis, as indicated.

### Stereotaxic injection of AAV2 vectors

Mice were anesthetized with isoflurane (1-5%) and placed in a stereotaxic frame (David Kopf Instruments). Injection was performed with a Wiretrol capillary micropipette (Drummond scientific) at a flow rate of 50 nL per minute (Micro4 controller, World Precision Instruments; Nanojector II, Drummond Scientific). The pipette was left on place for 10 minutes after the injection before retracting it slowly from the brain. *Alk4*^fl/fl^ mice were injected bilaterally with 300 nL of AAV2 adeno-associated viruses expressing either Cre recombinase and GFP (pAAV.CMV.HI.eGFP-Cre.WPRE.SV40) or only GFP (pAAV.CMV.PI.EGFP.WPRE.bGH) (Penn Vector Core, 2.27×10^13^ genomic copies per mL) using coordinates obtained from Paxinos and Franklin atlas as follows: AP +1.7 mm, mediolateral ML ±0.6 mm, dorsoventral DV –4.5 mm). To evaluate the infection efficiency of the vectors, RNA was extracted and Acvr1b mRNA (encoding ALK4) was measured by RT-qPCR (Figure S4F, G).

### Immunohistochemistry, in situ hybridization and image analysis

Mice were deeply anaesthetized with isoflurane (Baxter Medical AB) and perfused transcardially with 25ml of 0.125 M PBS (pH 7.4, Gibco) and 40 ml of 4% paraformaldehyde (PFA, Histolab Products AB). Brains were removed, post fixed in PFA overnight and cryoprotected in 30% sucrose in PBS. 30 μm brain sections were cut in a microtome (Leica SM2000 R). Sections were stored at -20 °C in a cryoprotective solution containing 1% DMSO (Sigma-Aldrich) and 20% glycerol (Sigma-Aldrich) in 0.05 M Tris-HCl pH 7.4 (Sigma-Aldrich). For RNA in situ hybridization, washing and fixation solutions were prepared using RNAase free components; surgical instruments and microtome blades were cleaned with RNaseZap (Sigma-Aldrich) before use.

For immunohistochemistry, brain sections were washed in PBS and blocked for 1 hour in 5% normal donkey serum (NDS, Jackson Immunoresearch) and 0.3% Triton X-100 (Sigma-Aldrich) in PBS. Incubation with primary antibodies was done overnight at 4 °C in blocking solution. After three consecutive 10 minute washes with PBS, sections were incubated with the corresponding secondary donkey Alexa fluorescent antibodies (1:1000, ThermoFisher) and 0.1 mg/ml of 40-6-diamidino-2-phenylindole (DAPI; Sigma-Aldrich) for 2 hours at room temperature. Sections were finally washed with PBS (3×10 minutes), mounted on glass slides in 0.2% solution of gelatin (Sigma-Aldrich) in 0.05 M Tris-HCl buffer (pH 7.4), dried and mounted with glass coverslips using DAKO fluorescent mounting medium. The following primary antibodies were used: rabbit anti ALK4 (1:500; ab109300, Abcam); mouse anti DAPP32 (1:750, sc-271111, Santa Cruz Biotech); rabbit anti DARPP-32 (1:500, sc-11365, Santa Cruz Biotech); mouse anti D1 (1:500; sc-33660, Santa Cruz Biotech); goat anti Activin A (1:250, AF338, RnD); mouse anti GFAP-Cy3 conjugated (1:1000, C9205, Sigma-Aldrich) and rabbit anti Iba1 (1:1000, Wako, 0719-19741).

For detection of Acvr1b mRNA by in situ hybridization, a template for riboprobe generation was obtained by RT-PCR from P1 mouse brain tissue using sense primer, 5′-TTTAAGCTGTTCCTCTGCCTAC -3′ and antisense primer, 5′-CCCAAGACTTCCACCTACATTC -3′ (Tm = 62°C) and subsequently subcloned into PCR II-TOPO-TA cloning vector (Invitrogen). Riboprobes were generated with a BIOT-NTP labeling kit (Roche). For the antisense riboprobe, the template plasmid was linearized with Spe1 (New England Biolabs) and transcribed with T7 polymerase (Roche); for the sense riboprobe, template was linearized with Xho1 (New England Biolabs) and transcribed with Sp6 polymerase (Roche). Coronal brain sections of 30 μm were obtained on a cryostat. Free-floating sections were washed in PBS followed by 5X saline-sodium citrate buffer (SCC) (0.75 M NaCl and 75 mM sodium citrate). The sections were then pre-hybridized for 2 hr at 50°C in SCC buffer containing 50% deionized formamide and 40 μg/mL salmon DNA (Thermofisher) and then incubated with sense or antisense probes at 75°C for 10 min, followed by 58°C for 16 hr. After washing at 65°C for 2 hours, the sections were treated with distilled water to quench endogenous autofluorescence, blocked in 0.5% tyramide signal amplification solution (TSA; TSA biotin kit; PerkinElmer) for 30 min, and stained with Streptavidin-HRP (1:100) at room temperature. After several washes, the sections were incubated in biotin-tyramide (1:100) reconstituted in amplification diluent (TSA biotin kit; PerkinElmer) for 10 min. After further washes, the sections were stained with Streptavidin-488 (Life Technologies; S32355; 1:1,500 dilution) for 90 min. Sections were then washed and prepared for immunohistochemistry as described above. Control hybridizations with sense riboprobe did not give any signal.

All fluorescence images were captured with a Carl Zeiss LSM 710 confocal microscope using ZEN2012 software (Carl Zeiss). Image analysis was performed using ImageJ and Fiji software packages. Quantification of the number PLA dots was done in imageJ using the plugin Synaptic Puncta Analyzer (Ippolito and Eroglu, 2010). Background was subtracted from the image and the same intensity threshold and range of particle size was used to analyze all images from the same experiment.

### Statistical analysis

Statistics analyses were performed using Prism 9 and Graph Plot for macOS(GraphPad Software, LLC.). Data are expressed as average ± S.E.M. Sample sizes were determined a priori as N=3-4 for in vitro studies and N=5-6 for in vivo studies based on previous work from our laboratory and other studies in the field. F-test or Kolmogorov-Smirnov test with Dallal-Wilkinson-Lillie for p value were used to assess data normality and homogeneity variance. Group comparisons were made using unpaired Student’s t-test, one-way ANOVA or two-way ANOVA as appropriate, followed by Bonferroni or Tukey’s multiple comparison tests for normally distributed data. Differences were considered significant when p< 0.05.

## Acknowledgements

The authors would like to thank Iina-Lotta Eleonoora Korkala (Karolinska Institute) for help with RT-qPCR; Linda Thors, Emma Wallet and Jessica Sundell (KMB animal facility, Karolinska Institute) for assistance with animal care; Albert Blanchart Aguado (VirusTech Core, Karolinska Institute) for assistance with production of shPCBP1-lentivirus; Klas Kullander (Uppsala University, Sweden) for *Gpr101*^Cre^ mice and all CIBLab members for comments and suggestions. Support for this research was provided by grants to C.F.I. from the Swedish Research Council (2016-01538 and 2020-01923).

## Author contributions

F.K. performed the majority of experimental work; D.F.S. contributed to experimental design, stereotaxic injection and in situ hybridization; A.A. performed all mouse genotyping and assisted in cell culture and histology studies; A.C.-R. performed the ALK4 immunohistochemistry. F.K. and C.F.I. designed the experiments; C.F.I. wrote the paper.

